# Palliative Irradiation Affects The Degradation Of Vertebral Bone Mechanics, Architecture, and Composition: A Longitudinal In Vivo Study In A Rat Model

**DOI:** 10.64898/2026.07.14.738210

**Authors:** Chenghua Wang, Madison Berardi, Samantha Martin, Christine Brown, Zahra Soltani, Mario Keko, Megan E. Rosa-Caldwell, Marie Mortreux, Seward B. Rutkove, Stacyann Bailey, Ron N. Alkalay

## Abstract

**Background:** Palliative radiation therapy (RT) for metastatic spine disease significantly increases the risk of vertebral fractures. However, the temporal mechanisms underlying radiation-induced vertebral bone fragility remain poorly understood.

**Objective:** To evaluate the longitudinal effects of a single high-dose irradiation, simulating palliative RT, on vertebral bone mechanical, architectural, and compositional properties in a healthy, skeletally mature rat model.

**Methods:** Thirty-one male Sprague Dawley rats received a single 15 Gy lumbar spine irradiation (IR). L4 vertebrae were assessed across all groups (irradiation: 7, 14, and 28 days post-IR, controls: at 0 and 28 days post-IR) for compressive strength and stiffness, micro-CT-derived bone composition and trabecular indices, serum bone turnover markers (NTX and BAP) and advanced glycation endproducts (AGEs).

**Results:** Irradiation induced progressive deterioration of vertebral bone mechanical properties, with strength decreasing up to 44% and stiffness up to 38% by 28 days post-IR, compared to 0- day controls. Trabecular bone exhibited reduced BMD, BV/TV, and Tb.N with increased Tb.Sp, a shift toward a more rod-like structure. Early post-IR changes suggested disrupted bone remodeling, characterized by elevated NTX and AGEs, but decreased BAP. Multivariable regression demonstrated that Tb.Th and AGEs were independent predictors of stiffness, collectively explaining 61% of its variance.

**Discussion:** High-dose irradiation induces sustained temporal degradation of vertebral mechanical properties driven by both trabecular architectural deterioration and alterations in bone matrix quality. Measures of bone composition and non-enzymatic bone turnover suggest this early damage was driven by disruption of bone cellular homeostasis, favoring increased resorption over formation. These findings support that radiation impairs both structural integrity and pre-yield mechanical behavior, providing mechanistic insight into the elevated fracture risk observed clinically after irradiation for metastatic spine disease.

**Lay summary:** This study used a rat model to mimic palliative radiation therapy for cancer that has spread to the spine and evaluated the changes in bone quality up to 28 days post-therapy. We found that irradiation progressively weakened the structural integrity and composition of the bones in the spine and disrupted the normal balance of bone breakdown and repair, leading to greater bone loss and fragility. Our findings provide insight into the increased risk of fractures observed in patients receiving radiation therapy to the spine and may support efforts to better protect bone health during treatment.

## 1. INTRODUCTION

Approximately 50% of all cancer patients will receive radiation therapy (RT) during their course of illness ^1^. In patients with metastatic spine disease (MSD), RT is routinely used to achieve local control of metastatic lesions and to help relieve pain ^1^. Palliative radiation therapy administered as either single-dose or multi-fractionation schemes over time, regardless of primary tumor histology, is thought to enhance tumoricidal effects by inducing apoptosis ^2^. RT increases the risk of vertebral fracture (VF) ^3^, reported to be 3% to 5% following conventional full-body irradiation^4^. Stereotactic body radiotherapy (SBRT), delivering ablative tumor doses while differentially sparing the adjacent spinal cord to a lower dose exposure, is associated with higher VF incidence rates, 13.9% and 39.0%, within six months following SBRT ^5^. The resulting higher risk is likely due to significantly higher deposition of radiation at the vertebral body ^5^. The initial fracture and any subsequent collapse can lead to catastrophic complications, including debilitating pain, spinal cord and nerve root compression, and paralysis ^6^. Clinical imaging-based diagnostics, reliant on either CT-based bone density assessment or score-based classification systems, have shown low sensitivity in predicting which patients will develop late-onset fragility ^7^. Although RT has been shown to lead to a reduction in vertebral bone mineral density (BMD) ^8,9^, limited clinical evidence exists to support the association between RT-induced BMD reduction and higher risk of VF ^9^. Hence, the underlying pathophysiology of radiation-induced bone damage on the risk of vertebral failure in MSD patients is not well understood.

Preclinical models have been essential for investigating the effects of radiation on bone quality and mechanical properties at different hierarchical bone structure levels ^10^. In hindlimb irradiation models, irradiation degrades bone vasculature ^11^, reduces osteocyte viability, increases osteoclast activity, and causes modification of the collagen structure and mineral chemistry ^10,12–14^. At the tissue level, irradiation causes higher accumulation of advanced glycation end product (AGEs) and mineral and matrix misalignment ^12,14–18^. At the whole bone level, single point and longitudinal assessment in tibial and femoral bones found irradiation decreased bone mass and resistance to fracture ^19,20^.

By comparison, few studies have investigated the specific effect of irradiation on vertebral bone. In preclinical rat spine models, application of fractionated (10 × 3 Gy) ^21^ or a single dose (20 Gy) ^22^ irradiation resulted in significant loss of vertebral strength at twelve (28%) and two (22%) weeks, respectively. Both studies demonstrated irradiation degrades vertebral bone architecture, although its effect on bone mineral density showed conflicting patterns. Furthermore, irradiation caused degradation of bone tissue composition indicated by elevated non-enzymatic collagen crosslinks ^21^ with the changes in the bone mass, architecture, and material properties accounting for 56%, 20%, and 24%, respectively, of the measured reduction in vertebral strength ^21^.

This study employed a Sprague Dawley rat model to characterize the effects of high-dose irradiation, simulating palliative treatment, on longitudinal changes in the mechanical properties of lumbar vertebral bone. Specifically, at each study timepoint following irradiation, we aimed to evaluate the relationship among radiation-induced changes in cancellous bone architecture, measured by high-resolution micro-computed tomography (micro-CT), tissue composition quantified using biochemical assays, and alterations in bone mechanical properties. To better understand these effects at the tissue level, we evaluated alterations in bone turnover through assessment of serum bone turnover markers (N-terminal telopeptide of type I collagen (NTX) and Bone Alkaline Phosphatase (BAP)) and bone matrix quality (AGEs). We hypothesized that high-dose irradiation would induce progressive, time-dependent deterioration of vertebral bone architecture, disrupt bone turnover, and reduce vertebral mechanical strength and stiffness in healthy Sprague–Dawley rats.

## 2. MATERIALS AND METHODS

### 2.1 Animals and Experimental Design

Beth Israel Deaconess Medical Center (BIDMC) Institutional Animal Care and Use Committee (IACUC) reviewed and approved animal experiments. Thirty-nine, 15-week-old male Sprague Dawley rats were obtained from Charles River Laboratories (Shrewsbury, MA, USA) and housed in the animal facility at BIDMC. Two rats were maintained per cage, with all animals allowed to acclimate for four days prior to the study. Water and food were provided ad libitum using an automatic serving system on a day/night cycle every 12 hours. The animals were randomly allocated at baseline to irradiated (IR) animals or control (C). Animals allocated to the irradiated group were euthanized post-irradiation (IR) at 7 days (7-IR: n=11), 14 days (14-IR:n=10) and 28 days (28-IR:n=10), Figure 1. The control group was euthanized at 0-day (0d-C) and 28-day (28d-C), n=4 each, Figure 1. Each animal’s body weight was acquired using a standard electronic scale at 0-day and on the day euthanized.

**Figure 1.**
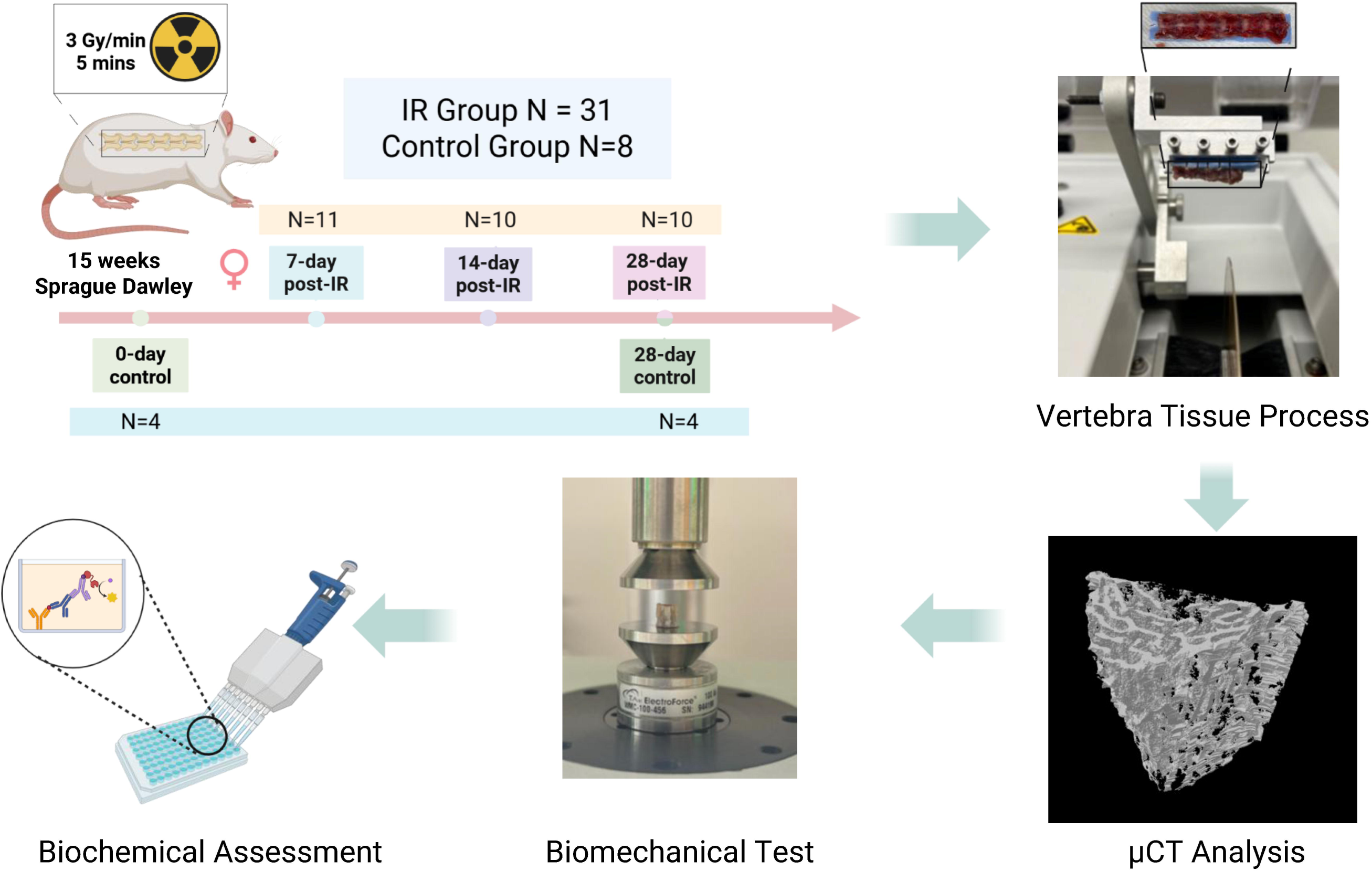
A graphical abstract of the study’s experimental design and analysis

### 2.2 Irradiation Protocol

Each animal was anesthetized via an intraperitoneal injection of ketamine 80 mg/kg and xylazine 5 mg/kg, with the required dose (0.1ml/100g) computed based on the animal’s weight. Ophthalmic ointment was applied to both eyes to prevent desiccation.

#### A. Irradiation experiment

Two rats were placed sideways back-to-back within the enclosure of the animal irradiator (X-Rad32 Precision X-Ray Inc, Madison, CT, USA). A custom-made 6.5mm lead shield, designed to limit exposure only to the lumbar spine area, was placed on each animal (Figure 2). Using Six nanoDot™ dosimetry badges (Landauer, Glenwood, IL, USA), a pilot measurement to measure the shield effectiveness determined the lead shield attenuation to follow an exponential relationship (Log(Gy) = 2.635 - 1.033 × lead thickness(mm). Based on these measurements, the 6.5 mm shielding transmitted radiation amounted to <0.1Gy.

**Figure 2.**
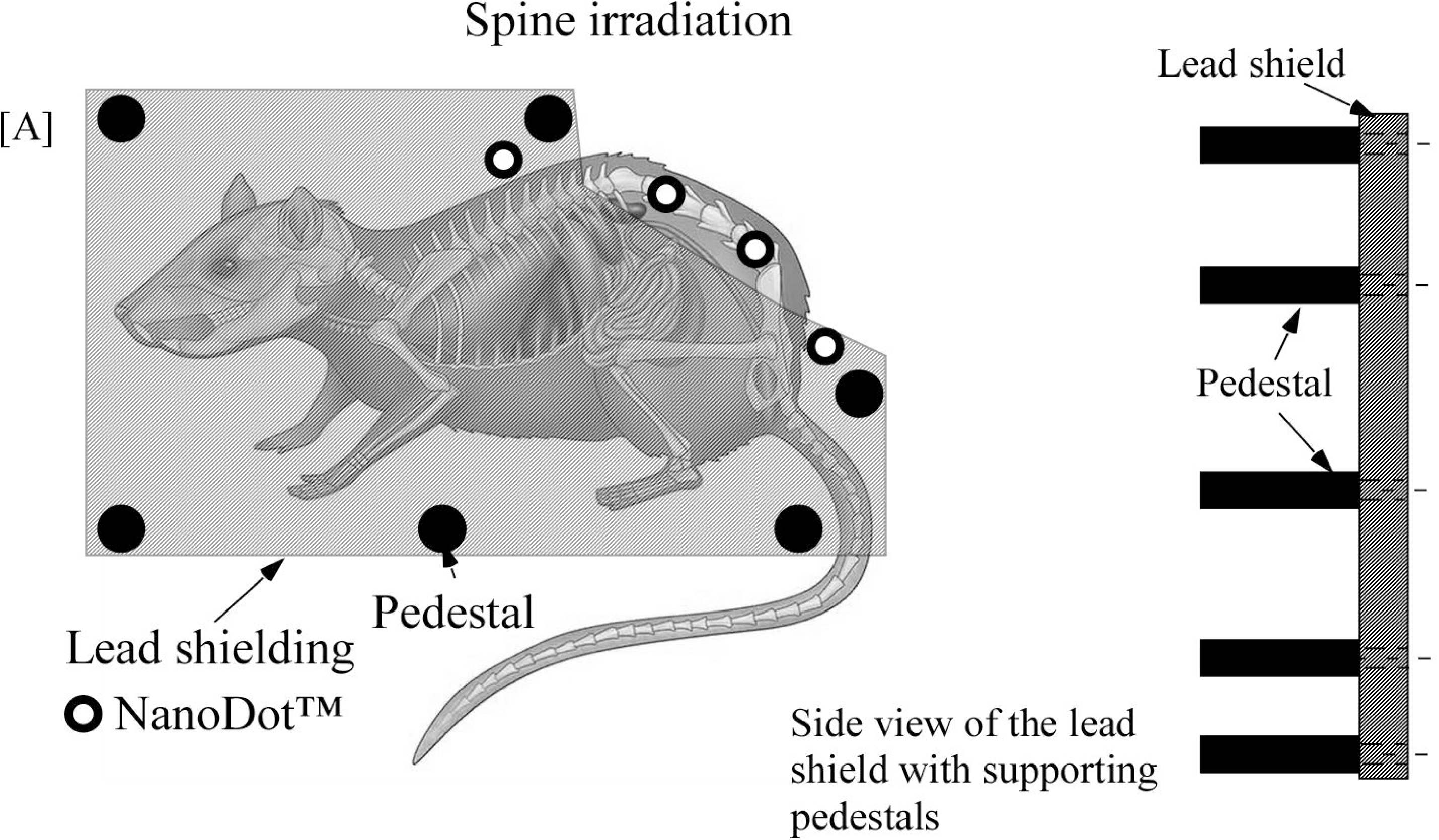
Graphic illustration of the irradiation experiment setup, illustrating the position of the rat and NanoDots™ under the lead shield.

For the experiment, six nanoDot™ dosimetry badges (Landauer, Glenwood, IL, USA) were placed next to the irradiated spine and on the animal’s body within the shield (Figure 2) to measure the applied radiation and the shield’s effectiveness. With the shield and nanoDot™ situated, the animals were irradiated with 15Gy at a rate of 3Gy/min for five minutes (320KV, 12.5mA, 50cm SSD (HVL≈ 1mm Cu). Assuming a normal tissue alpha/beta of 3.0, the biologically equivalent dose (BED) of 15 Gy in 1 fraction is 15 × (1+15/1/3)= 90.0 Gy. This dose would be the equivalent of 30 Gy in 5 fractions, for which the BED would also be 30 × (1+30/5/3) = 90 Gy. This dose is within the range of 20-30 Gy in 5 fractions that would often be given for palliation by conventional or stereotactic RT ^23^.

#### B. Post-irradiation procedure

Upon the completion of the irradiation experiment (total time =15 minutes), the animals were transferred to a microisolator cage with a water heating pad during recovery (@ 37° C, TP500 T/Pump, Gaymar Industries, Inc., Orchard Park, NY, USA) under supervision until ambulatory. Once recovered, the animals were transferred back to the animal facility. Each animal was monitored daily for five days for signs of pain, distress, sepsis, or altered food and water intake. Based on the animal’s Body Condition Score (BCS) and/or weight loss, the veterinarian was consulted if an animal required veterinary care. None of the animals required such care.

### 2.3 Specimen retrieval and preparation

All animals were euthanized using a SMARTBOX™ CO2 chamber System (E-Z Systems Inc., Bethlehem, PA, USA) with secondary confirmation of the euthanasia performed using a thoracic puncture.

#### A. Blood sample collection

At the time of thoracic (cardiac) puncture, we used a 23G needle to withdraw 10ml of blood. The blood was collected in a red-top tube, and it was allowed to clot undisturbed for 30 minutes at room temperature, followed by centrifugation for 10 minutes to separate its components. Serum was then separated, transferred to a new collection tube and stored at -80°C.

#### B. Spine segment collection

The lumbar spine L1-L6 with surrounding spinal musculature was surgically extracted en bloc, and the segment was cleaned of all muscle, maintaining the ligamentous tissues and the intervening as well as the intervertebral discs adjacent to L1 and L6. The spine was extracted, cleaned of oil, wrapped in saline-soaked gauze, and stored at -20 ° C in a double plastic bag.

### 2.4 MicroCT imaging-based assessment of bone architecture

#### A. Retrieval of L4 level

On the day of imaging, each spine was thawed at 4°C (12 hours), placed in a custom fixation device, and the posterior elements were embedded in dental cement such that the vertebral bodies and pedicles were exposed for sectioning. The device was rigidly positioned in a gravity-fed precision sectioning system (IsoMet 1000, Buehler, IL, USA), and the diamond-coated circular saw was used to obtain a plane-parallel L4 level under constant irrigation.

#### b. MicroCT imaging

Each vertebral section was scanned with μCT (SKYSCAN 1276, Bruker Corp., Billerica, MA, USA) using the following parameters: 85 kV, 200 µA, 10.2 µm voxel size, 684 ms exposure time, 360° rotation, 0.4° rotation step, frame averaging x2, and 1mm Al filter. The data was reconstructed using the NRecon software (Bruker Corp., Billerica, MA, USA) with the ring artifacts reduction factor set at 7, beam hardening correction set at 30%, and dynamic range set at (0-0.038). A global threshold value of 423 mgHA/cm^3^ was used for bone segmentation, and the μCT analysis software was used to compute the following bone architectural parameters: bone volume fraction (BV/TV), trabecular number (Tb.N), trabecular thickness (Tb.Th), trabecular separation (Tb.Sp), and connectivity density (Conn.D). Bone mineral density (mgHA/cc) was computed from the complete bone core volume segmentation.

### 2.5 Biomechanical Testing

The L4 vertebral level was positioned at the center of a compression test device secured to an ElectroForce 3200 Series III electro-mechanical test system (TA Instruments, New Castle, DE, USA). The test system was used to apply a pre-compressive load of 0.5N to ensure contact between the specimen and the compression platen, followed by a monotonic ramp compressive displacement at a rate of 1.0 mm/min until the vertebral failure was registered. This failure was defined as 5% loss of maximum compressive force recorded (TA Instruments, model 944199, 450N). The test system acquired the applied displacement and the resulting compressive load at a rate of 100Hz. At test completion, the vertebral sample was wrapped in saline-soaked gauze and stored at -20°C in a double plastic bag. The test load-displacement curve was analyzed to compute 1) vertebral strength, defined as maximum measured compressive force (N), and 2) vertebral stiffness, defined as the slope coefficient from a linear regression model fitted to 20-80% of the curve to max failure load.

### 2.6 Biochemical Markers of Bone Remodeling

#### A. Total Fluorescent Advanced Glycation End-products

Each mechanically tested vertebra was thawed, defatted using isopropyl ether, lyophilized (freeze-dried) overnight, and hydrolyzed in 6N hydrochloric acid (HCL) based on dry weight (31-47 mg dry weight) for 16 hours ^24^. Auto fluorescent protein crosslinks were quantified in each hydrolysate using a quinine sulfate standard (stock: 1 μg quinine per mL of 0.1N sulfuric acid) at 360/460 excitation/emission using an Enspire 2300 microplate reader (PerkinElmer, Waltham, MA, USA). The collagen content in each hydrolysate was calculated using a hydroxyproline standard of increasing concentration (stock: 2000 μg L-hydroxyproline per mL of 0.001 N HCl). The absorbance of the hydrolysates and the hydroxyproline standard was subsequently measured at 570nm. Total fluorescent AGEs were computed as the amount of quinine (ng) per amount of collagen (mg) ^24^.

#### B. Measurement of Serum Biomarkers

**Bone alkaline phosphatase (BAP)** was measured from the serum of all animals using a sandwich enzyme-linked immunosorbent assay (ELISA) kit from Lifeome Biolabs (Cat # ER0761-6, San Diego, CA, USA). Briefly, all samples, standards, and biotin-conjugated detection antibody were incubated on a 96-well plate pre-coated with a bone-specific ALP capture antibody. The plate was washed, then incubated with horseradish peroxidase (HRP)-streptavidin. A TMB substrate was used to visualize the HRP enzymatic reaction, and the absorbance was read at 450 nm.

**Crosslinked N-Telopeptide of Type 1 Collagen (NTX)** was measured from an aliquot of the serum using a competitive ELISA kit from Cloud-Clone Corp (Cat # CEA639Ra, Katy, TX). Briefly, a monoclonal antibody specific to NTX was pre-coated onto a 96-well plate. A competitive binding reaction occurred in which unlabeled NTX present in the samples or standards competed with biotin-labeled NTX for a limited number of antibody sites when incubated together. After incubation, unbound material was washed off, and avidin-horseradish peroxide (HRP) was added to all wells. The amount of HRP retained in the well is inversely proportional to the NTX concentration and, consequently, to the absorbance measured at 450 nm.

### 2.7 Statistical Analysis

We created box plots to graphically summarize the longitudinal effects of the irradiation experiment (7-IR, 14-IR and 28-IR) as well as the sham irradiation control groups (0d-C and 28d-C) on bone architecture, composition, remodeling biomarkers, and the vertebral mechanical properties. We used Welch ANOVA models to compare these parameters by group and to estimate the following post-hoc comparisons: 1) 7-IR, 14-IR, 28-IR vs. 0d-C and 2) at 28-IR vs. 28d-C. We included all the available data in the ANOVA models and verified the assumptions. We used the SAS software (SAS/STAT® version 9.4, SAS Institute Inc., 2013, Cary, NC) to implement the ANOVA models. We used the R software (version 4.3.1, R Core Team (2023), R Foundation for Statistical Computing, Vienna, Austria) to create graphical summaries and to calculate the Pearson correlation between the vertebral mechanical properties, bone architecture, composition and the remodeling biomarkers at animal termination, with the results presented as a correlation heatmap. For all the analyses, we controlled for multiplicity by implementing the False Discovery Rate approach ^25^. We used adaptive lasso regression to identify the most important predictors for strength and stiffness. We employed a separate model for each outcome and referred to the R2 to assess the proportion of variance explained by each model.

## 3. RESULTS

### 3.1 Palliative irradiation negatively affects Animals’ weights

The Welch ANOVA showed a significant difference in the weight of the animals between the study groups (p=0.0003), with the 7-IR group having significantly lower weight (15.5% lower) (p=0.0004) compared to the 0d-C. Weight was 4.6% and 3.7% lower in the 14-IR and 28-IR groups, respectively, but not significantly different from 0d-C. By contrast, at 28 days, the control group (28d-C) weight was higher than 0d-C, a difference of [7.9 %, (p=0.0491)], Figure 3.

**Figure 3.**
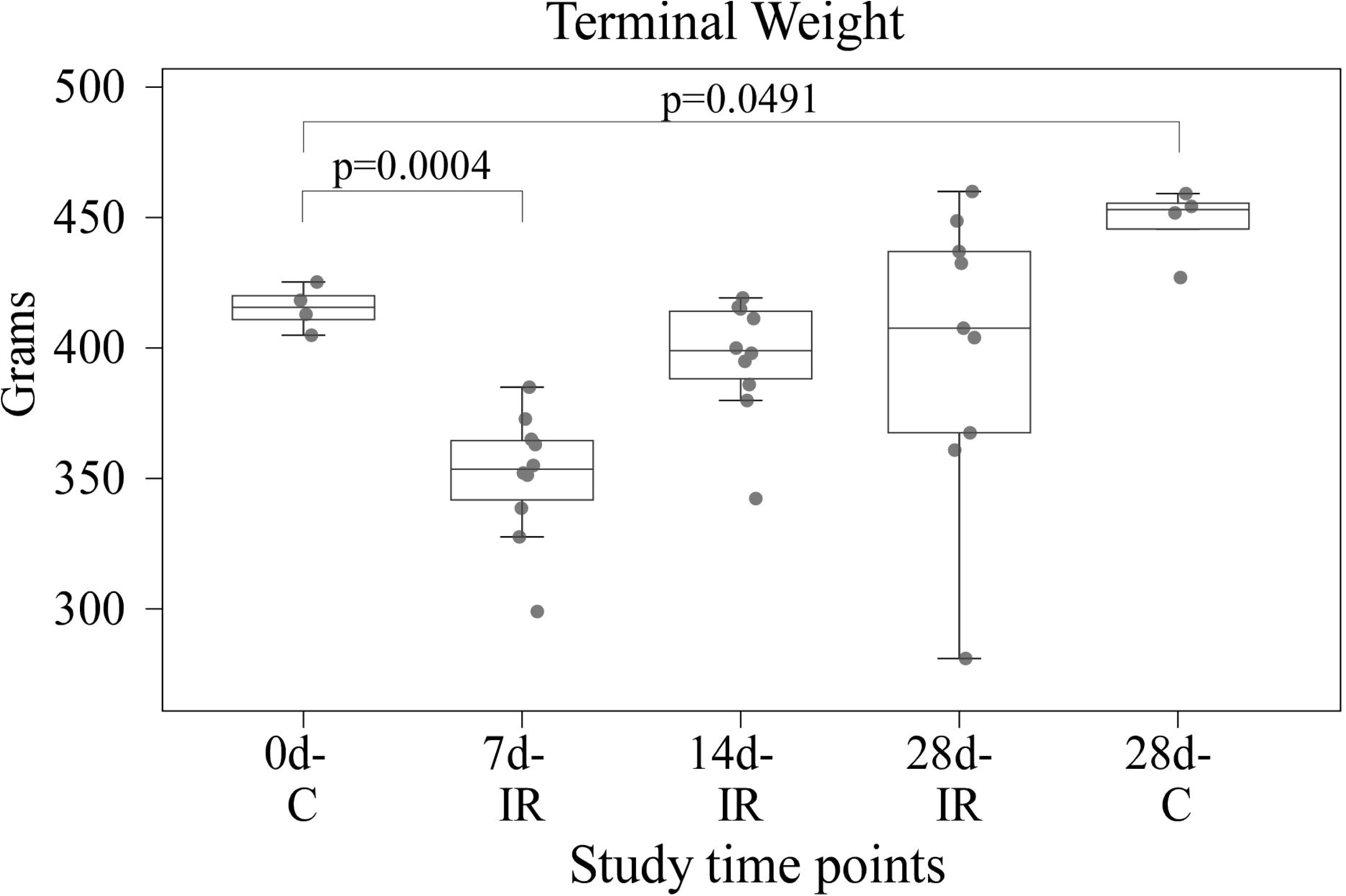
Statistical summary for the longitudinal change in animal weight for the irradiated and control animals at the experimental endpoints post-irradiation.

### 3.2 Irradiated vertebrae show temporal degradation of mechanical properties

The Welch ANOVA analysis revealed that irradiation had a significant and consistent negative effect on vertebral strength and stiffness as measured by mechanical testing (strength: p=0.0014, Stiffness: p=0.0003). Irradiation caused a near-monotonic decrement in the irradiated vertebrae strength, resulting in the mean strength value being lower by 22.6% at 7-IR and 32.0% at 14-IR compared to 0d-C. The difference was significant at 28-IR, with a decrease by 44.4% (p=0.0229), Figure 4. A similar pattern of decline was observed in measured stiffness: -1.1% at 7-IR and - 12.5% at 14-IR compared to 0d-C. The decrease was significant 28-IR, (-38.0%, p=0.0006), Figure 4. In comparison to the 28-IR group, the older control animals (28d-C) exhibited higher measured strength (168.2%, p=0.0006) and stiffness (90.2%, p<0.0001). Similarly, the 28d-C showed higher strength and stiffness than the 0d-C group (49.1%, p=0.0220 and 17.9%, p=0.0230), respectively (Figure 4).

**Figure 4.**
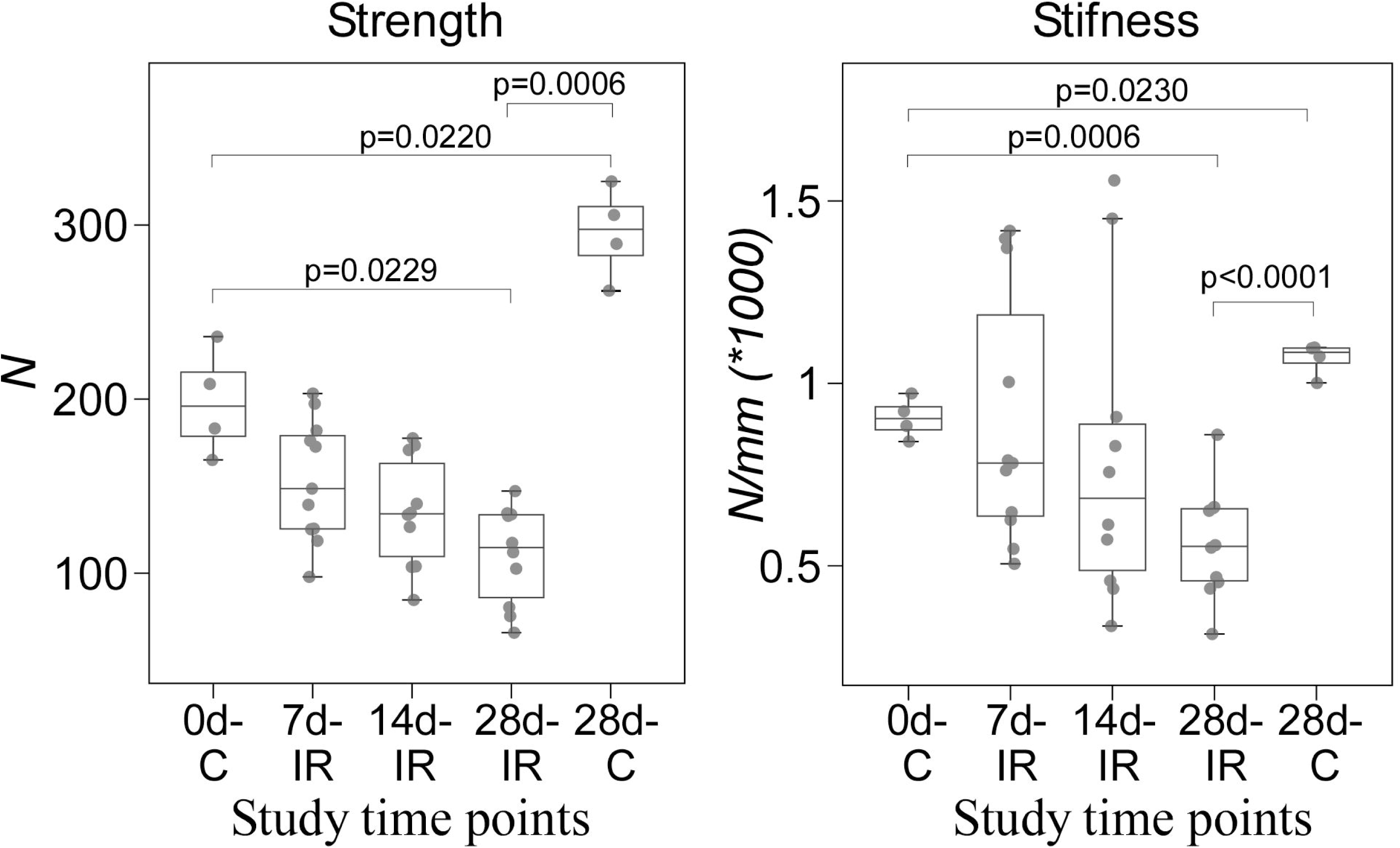
Compared to the control groups, irradiation caused a near-monotonic decrease in vertebral strength and stiffness.

### 3.3 Irradiation induces loss of vertebral bone density, relative volume and architecture over time

Compared to the 0d-C group, irradiation caused a moderate decrement in vertebral bone mass (BMD) and tissue volume fraction BV/TV) at 7-IR (-5.6% and -9.9%, respectively) and at 14-IR (- 8.4% and -13.9%, respectively). The decrease was significantly higher at 28-IR (-33.9%, p=0.0230 and -42.6%, p=0.0361, respectively), Figure 5.

**Figure 5:**
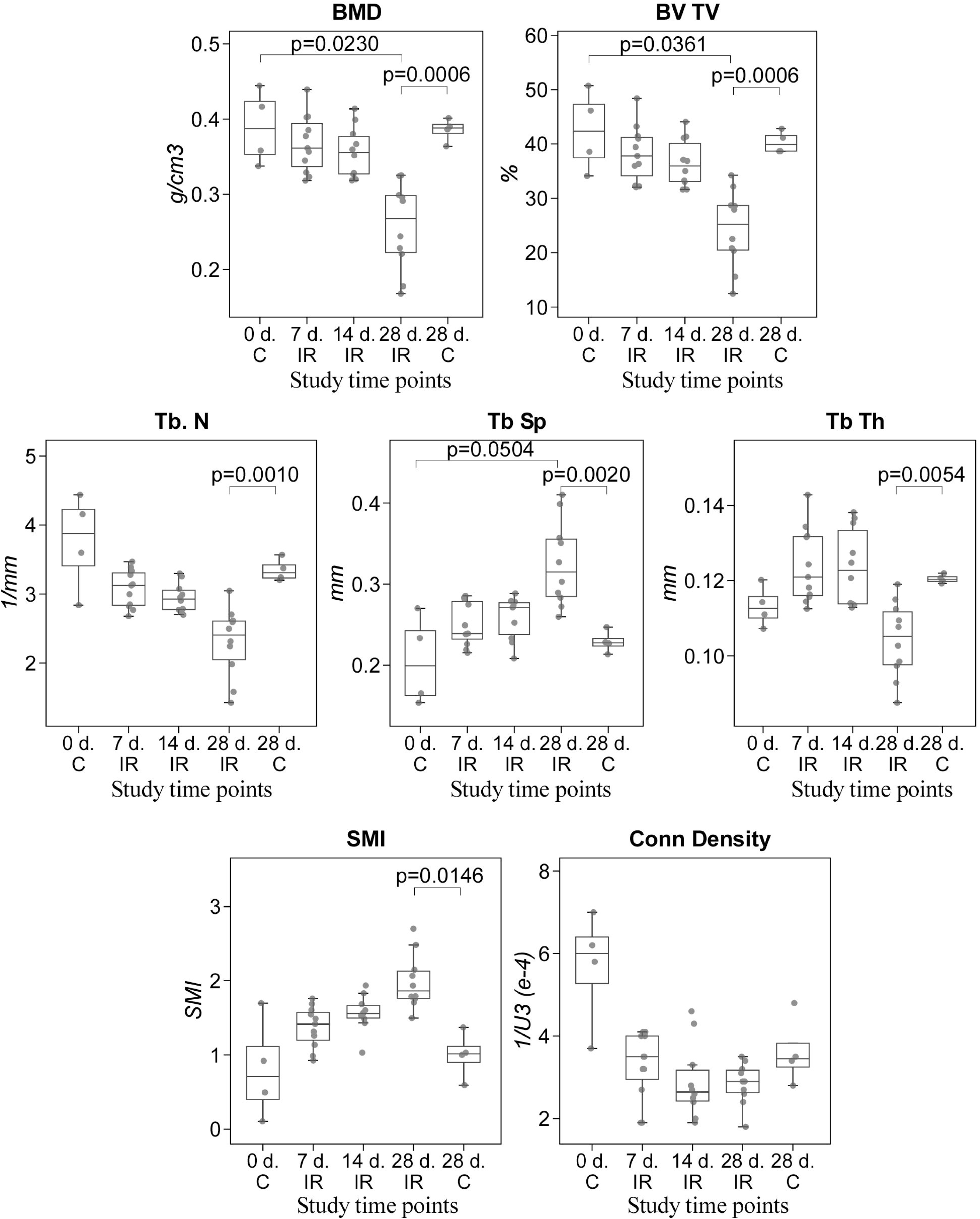
ANOVA analysis of the longitudinal changes in primary and secondary architectural indices demonstrates that irradiation affects degradation at early stages post-treatment, the degradation showing relative worsening at the later stage of the study. * (p<0.05) and ** (p<0.001) indicate statistically significant differences between the test group and 0-C. + (p<0.5) and ++ (p<0.001) indicate a statistically significant difference between 28-IR and 28d-C.

Compared to the 0d-C group, irradiation caused marked degradation in the trabecular bone architecture, manifested at 7-IR and 14-IR by lower mean Tb.N (7-IR: -18.1%; 14-IR: -21.6%) and Conn.D (7-IR: -42.2%; 14-IR: -48.8%), and higher mean Tb.Sp (7-IR: 21.6%,; 14-IR: 25.7%), mean Tb.Th (7-IR: 9.5%; 14-IR: 9.3%) and mean SMI (7-IR: 70.8%; 14-IR: 94.4%) values, Figure 5. These differences were not significant at the 5% level. This trend worsened in the 28-IR group, with a greater decrement for mean Tb.Sp (58.1%), mean Tb.N (-38.8%) and Tb.Th (-7.9%) but higher mean Tb.Sp (58.1%, p=0.0470) and mean SMI (146.8%) than 0d-C. Figure 6, presenting the microCT image data corresponding to the study experimental time group, illustrates these changes.

**Figure 6:**
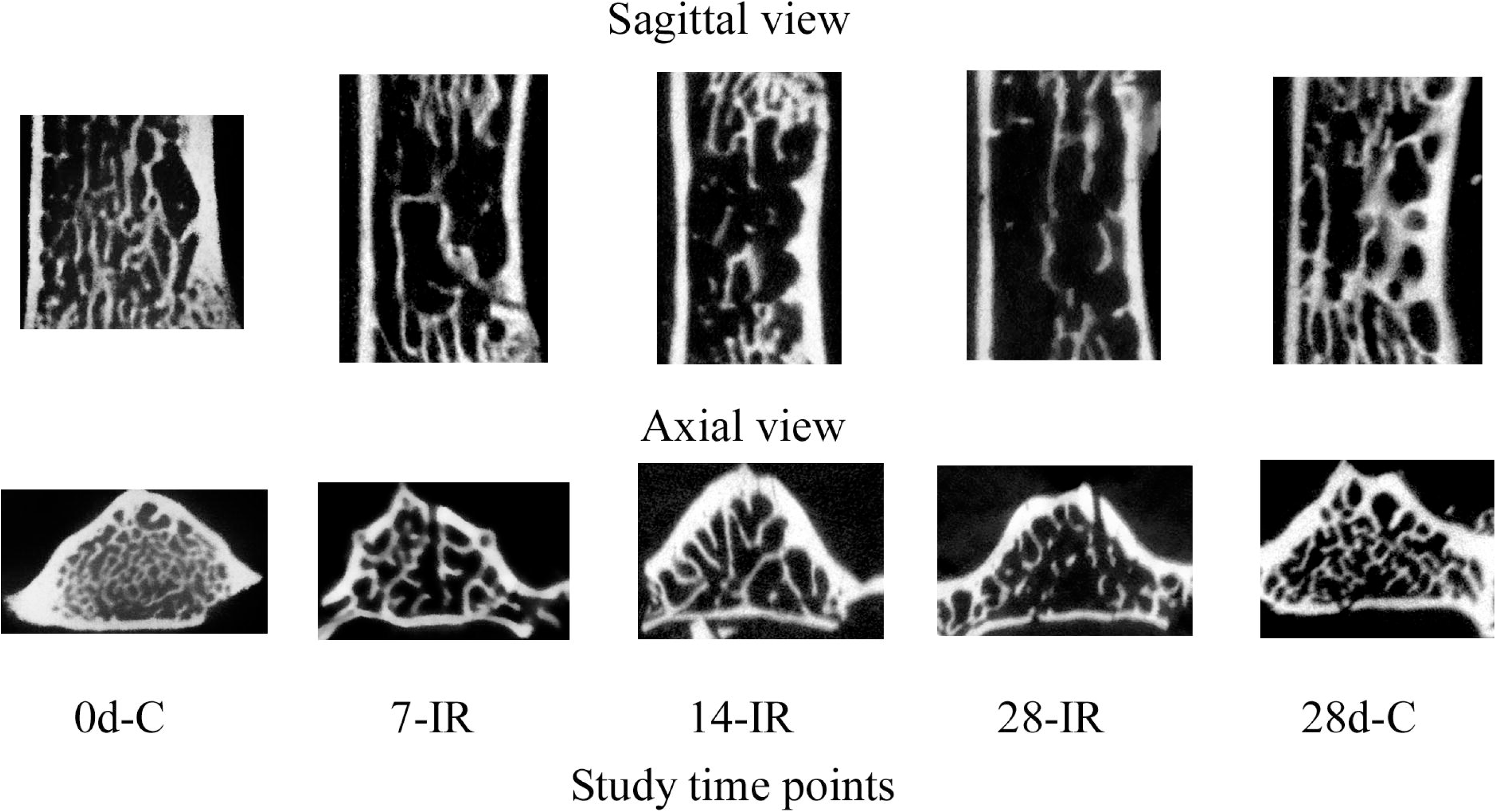
Sagittal and axial microCT data for vertebrae at the study time points demonstrate the early loss of bone architecture post-RT (7-IR), an apparent attempt at recovery at 14 days (14-IR), and the collapse of the repair response at 28 days (29-IR) post RT. In comparison to the 0d-C group, the 28d-C group showed maturation of the spatial bone structure.

Compared to the 28d-C group, the 28-IR group demonstrated significantly lower mean BMD (- 33.2%, p=0.0006), BV/TV (-39.6%, p=0.0006), Tb.N (-31.3%, p=0.0010), and Tb.Th (-13.5%, p=0.0054), and a significantly higher mean Tb.Sp (42.0%, p=0.0020) and SMI (98.9%, p=0.0146), Figure 6. Although mean Conn.D was markedly lower (-21.5%), this difference was not significant (p= 0.2340).

### 3.4 Irradiation affects time-dependent alteration in bone remodeling and composition

Figure 7 summarizes the post-irradiation temporal changes in TMD, collagen content, AGEs, NTX and BAP. Compared to 0d-C, the 7-IR group showed higher mean TMD (5.2%), NTX (41.6%), and AGEs (69.3%) and lower mean BAP (-28.3%). These differences were not significant at the 5% level. At 14-IR, this trend continued for mean TMD (7.1%, p=0.0446), NTX (46.3%), AGEs (18.0%) and BAP (-26.8%) while collagen content values showed little change compared to 0d-C. These differences were not significant at 5% level. Compared to 0d-C, the 28-IR group showed higher AGEs (46.5%), NTX (33.9%), collagen content (6.8%) and TMD (1.6%), with BAP values being lower (-33.2%). The differences for NTX, AGEs and BAP were not significant at the 5% level.

**Figure 7:**
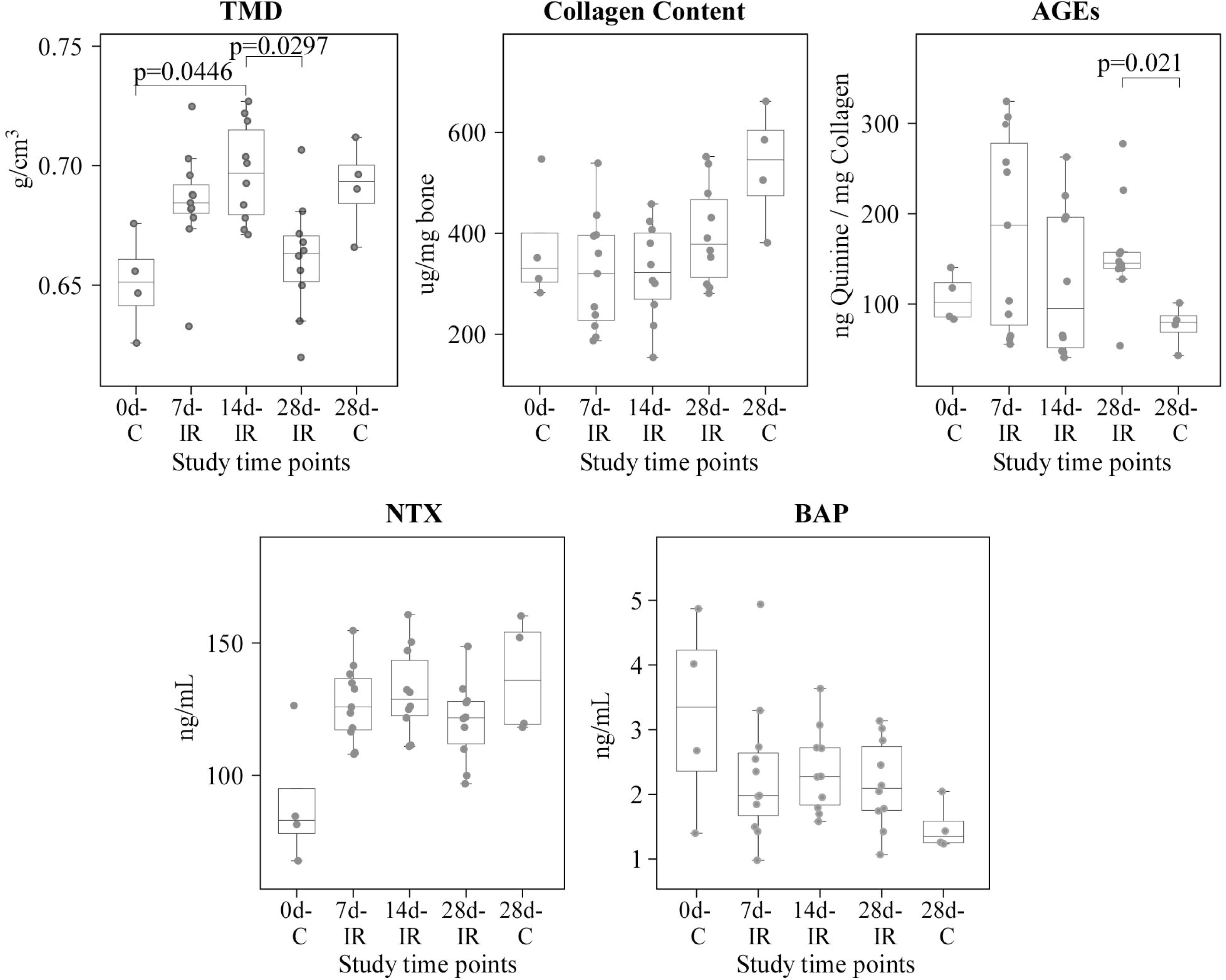
ANOVA analysis demonstrates irradiation affects early longitudinal changes in bone tissue composition (TMD, collagen content) with the changes in AGEs and bone turnover markers suggesting modification of the bone matrix quality and homeostasis.

Compared to the 28d-C, the 28-IR group showed significantly higher AGEs (106.4%, p=0.0209). Mean BAP values were markedly higher (44.9%), with mean collagen content (- 25.4%), NTX (-12.4%), and TMD (-4.3%) being lower. These differences were not significant at 5% level.

#### 3.5: Bivariate relationships between vertebral bone mechanics, architectural and compositional parameters at terminal assessment

Figure 8 presents a heatmap for the correlation between the irradiated vertebrae’s biomechanical properties, bone architectural indices, tissue composition, and turnover markers at terminal assessment. The corresponding heat map for the control groups is presented in the appendix, Figure A.1.

**Figure 8:**
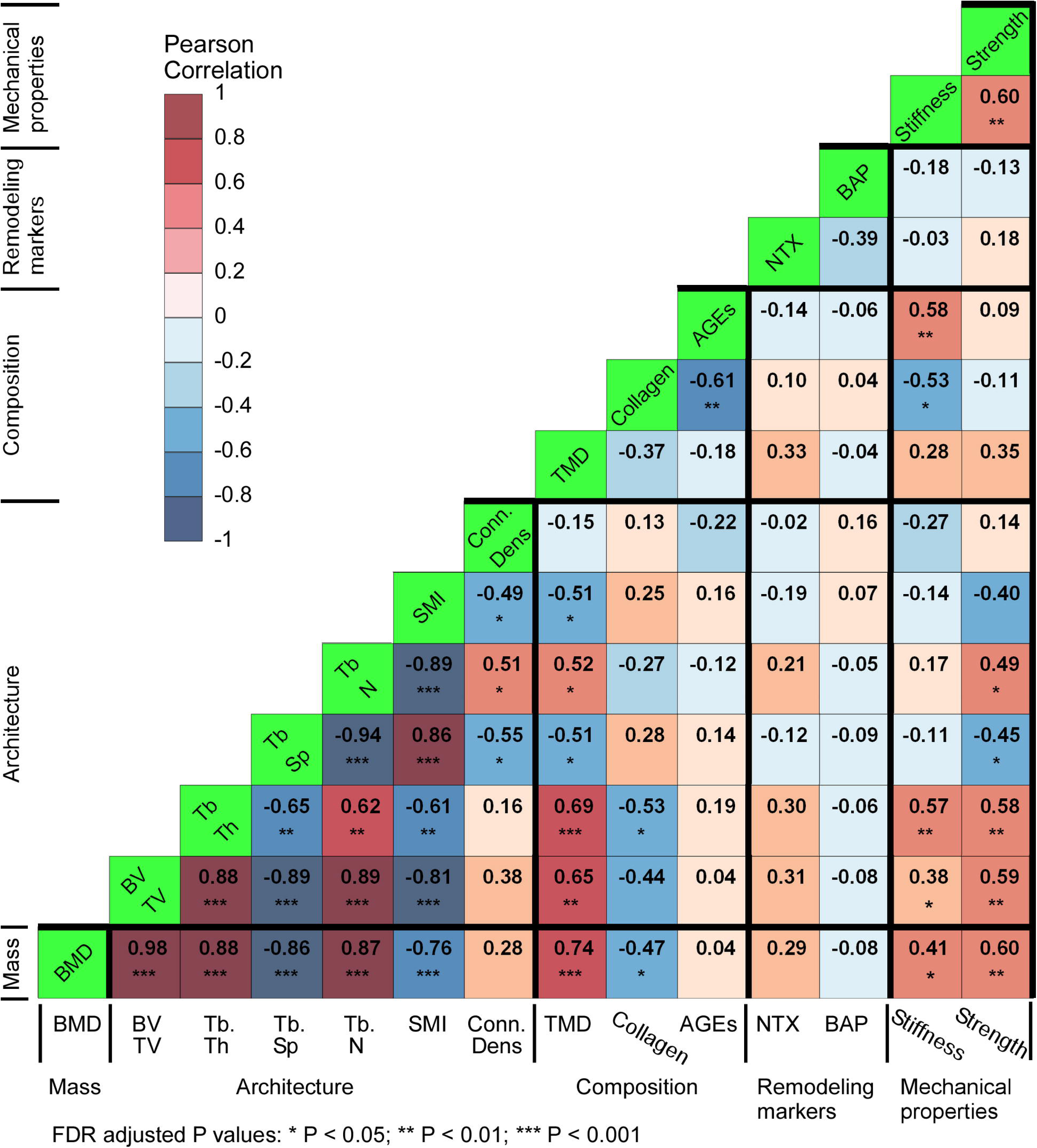
Heat map highlighting significant correlations between the bone architectural indices, composition and mechanical properties in the irradiated vertebrae at terminal assessment. Colors closer to brown indicate higher positive correlation, and colors closer to blue represent negative correlation.

#### a. Vertebral strength

The irradiated vertebrae’s strength correlated with higher vertebral stiffness (r=0.60, p=0.0003), bone mineral density (BMD: r=0.60, p=0.0004) and fraction (BV/TV: r=0.59, p=0.0005), and with thicker (Tb.Th: r=0.58, p=0.0007) less spaced (Tb.Sp: r=-0.45, p=0.0107), and more numerous (Tb.N: r=0.49, p=0.0047) bone trabeculae, Figure 8.

In the control vertebrae, strength correlated with higher stiffness (r=0.88, p=0.0039), collagen content (r=0.88, p=0.0039), and bone tissue density (TMD: r=0.81, p=0.0149), and with lower bone connectivity (Conn.D, r=-0.76, P=0.0260) and AGEs (r=-0.76, p=0.0280), Figure A.1.

#### b. Vertebral stiffness

In the irradiated vertebrae, higher estimated stiffness was correlated with thicker trabeculae (Tb.Th: r= 0.57, p=0.0008), and higher AGEs (r=0.58, p=0.0006), bone mass (BMD: r=0.41, p=0.0234) and fraction (BV/TV: r=0.38, p=0.0370), Figure 8, but lower collagen content (r=-0.53, p=0.0022). In the control vertebrae, (Figure A.1), higher estimated stiffness correlated with lower bone connectivity (Conn.D, r=-0.93, P=0.0009), and higher collagen content (r=0.74, p=0.0366), NTX (r=0.74, p=0.0366) and tissue mineral density (TMD: r=0.76, p=0.0280).

### 3.6 : Contribution of the vertebral bone parameters to its strength and stiffness

Derived from the adaptive lasso parameter selection, multivariable regression analysis Equation 1, found 76% of the variance in the measured strength explained by higher vertebral stiffness, BMD, and collagen content (p<0.0001). Independently, regression analysis revealed vertebral stiffness to explain 32% (p=0.002), BMD 28% (p=0.005) and collagen content 15% (p=0.0131) of the variance in strength, respectively.

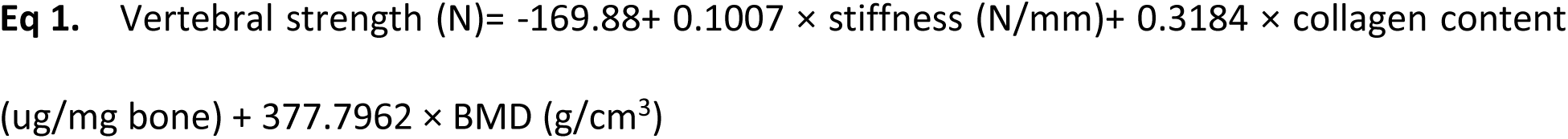

For vertebral stiffness, the multivariable regression model, Equation 2, revealed higher trabecular bone thickness and AGEs and lower collagen content and NTX explained 61% of the variance in the estimated stiffness (p<0.0001). Independently higher trabecular bone thickness explained 39% (p<0.0001) of the variance in vertebral stiffness, and AGEs explained 21% (p=0.0036) of this variance. Independently, the model found higher collagen content, explaining 7% of the variance, which was not statistically significant at the 5% level.

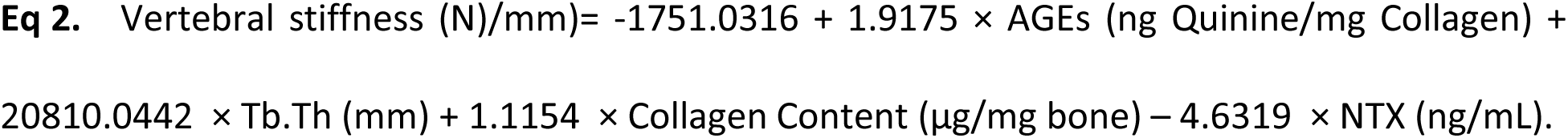

## 4. DISCUSSION

In a healthy rat model of high-dose palliative irradiation to the lumbar spine, we observed the early loss of vertebral body strength and stiffness with concomitant decrease in vertebral cancellous bone mass and volume fraction, the magnitude of loss becoming significant at the later stage of the study. The loss of vertebral stiffness, correlating with thinner bone trabeculae and lower AGEs, suggests that irradiation degrades the pre-yield properties of the vertebral bone via the alteration of bone collagen matrix and degradation of the bone trabeculae mechanical stability. Independently, the loss of vertebral strength beyond the physical reduction in bone mass suggests that high-dose irradiation disturbs the vertebral mechano-structural homeostasis. Careful consideration is needed in extending these findings to human patients receiving palliative radiation. Nevertheless, data provide a better understanding of how changes in bone microarchitecture and biochemical markers may underlie the clinically reported increased risk of vertebral fracture in cancer patients within a year after being treated with irradiation therapy.

### 4.1 Effect of irradiation on vertebral bone architecture and composition

Preclinical irradiation spine models demonstrated that low radiation doses, <= 5 Gy, cause a decrease in trabecular number and increased trabecular spacing, causing degradation of bone architecture ^21,26,27^. At the tissue level, single-dose focal IR affected persistent fragility, although tissue-level mineralization changes were less pronounced ^21,28,29^. Longitudinal studies found irradiation to cause early increases in trabecular spacing and higher SMI, but only modest changes in trabecular thickness, with the change in mineral density metrics lagging or remaining subtle initially ^27,30^. Together, these studies show the trabecular matrix is sensitive to irradiation. We employed 15-week-old rats upon entering the study, corresponding to late adolescence, and reached 19 weeks by the endpoint, which is considered a mature age ^31^. At this stage, physiological bone turnover is naturally slowing as growth nears completion ^32^. Ideally, this is reflected in the 28-day control (28D-C) group, which showed significantly lower BAP and higher NTX levels compared to the 0-day control (0D-C), indicating decreased bone formation and increased resorption due to maturation ^33^. The single high-dose irradiation experiment caused marked loss of bone mass and fraction at the study’s early stage (7-IR), with the loss worsening in the latter period of the study (14-28 days IR). Analysis of the micro-CT data showed this pattern of loss correlated with the increase in trabecular spacing, the decrease in bone trabecular number, the thinning of the bone trabeculae, and an increase in trabecular spacing, accompanied by loss of bone connectivity. The apparent degradation in bone architecture parameters was significant for all parameters analyzed compared to the 28-day control group. In agreement with previous murine studies using fractionated IR ^21,22,26,30,34^, our findings suggest irradiation affects a longitudinal-based rarefaction of the spatial vertebral bone architecture via a shift toward a more rod-like structure with the trabeculae undergoing perforation, as indicated by image-based bone topology measures (SMI) ^27,35^.

Reflecting on the observed architectural changes, the biochemical assay suggests that irradiation disrupted the bone turnover process at an early stage post-treatment. In comparison to the 0-Day controls, we observed an early, sharp decline in BAP to levels comparable with the 28d-C group at day 7-IR. Concomitantly, NTX levels in the IR group were markedly higher at 7-IR, indicating accelerated bone resorption ^14,36^. Although these changes were not significant, the data suggest that osteoblast activity was severely impaired ^13,16^, affecting the repair response of the bone. AGEs values are an indicator of irreversible, non-enzymatic collagen crosslinks that accumulate in bone type I collagen, especially as pentosidine, carboxy-methyl-lysine(CML), and *Nɛ*-carboxy-ethyl-lysine(CEL) ^24^. Specifically, impaired osteocyte function post-irradiation exposure could decrease the mechanosensitivity of bone ^14^, leading to impaired bone homeostasis resulting in alterations to bone mass and material (TMD). The current study’s observation of elevated levels of AGEs likely reflects higher heterogeneity in collagen crosslinking due to radiation-induced oxidation ^24^. The high variability in AGEs at 7 and 14 days indicates uneven bone remodeling, further supporting the indications regarding radiation-induced impairment of bone homeostasis, which aligns with the significant architectural changes observed by day 28, as previously discussed ^37^.

Previous studies showed irradiation doses in the 10–20 Gy range can induce pronounced osteocyte senescence, osteocyte apoptosis with elevated RANKL and sclerostin levels, which favor resorption over formation ^15,18,38^, and compromised lacunar–canalicular networks ^16,26,37^. These cellular-level changes were accompanied by altered bone composition and bone structural deterioration ^39,40^. Notably, comprehensive reviews underscore that doses exceeding ∼5 Gy suppress osteoblast proliferation and trigger osteocyte death, while multilevel assessments affirm that higher doses significantly deteriorate microarchitecture ^41^. Together, these prior findings may explain the decline in trabecular bone indices observed in our irradiation groups.

### 4.2 Effect of irradiation on vertebral structural response

In agreement with previous preclinical mouse and rat irradiation studies ^12,16,17,21,22^, our correlation analysis of irradiated animals demonstrated an association between loss of vertebral strength and lower bone mass and fraction, and degradation in vertebral bone architecture, shown by thinning of bone trabeculae with higher intertrabecular spaces and a lower number. This pattern is in marked contrast to the values exhibited by the mature sham irradiated controls, suggesting irradiation interferes with the process of bone remodeling, which occurs during the natural process of skeletal maturation. Significantly, regression analysis demonstrated vertebral stiffness to be a stronger determinant of vertebral strength compared to vertebral bone mass, suggesting irradiation has a concurrent effect on the vertebral mechanical properties. The dominant role of stiffness could be related to a few mechanisms.

Simulating the effect of irradiation on the state of stress within murine vertebrae, finite element (FE) modeling indicated irradiation-mediated reduction in Tb.N caused increased stress within the cancellous bone compartment ^21,29^. Furthermore, Alwood et al. ^29^ highlighted irradiation to reduce the bone modulus, resulting in the redistribution of loading and thus stresses towards the vertebral cortex ^29^. In this study, sham-irradiated vertebral strength and stiffness were associated with thicker bone trabeculae, lower trabecular number and bone connectivity, indicating a more plate-based structure. By contrast, the irradiated vertebrae strength was associated with the thinning of the bone trabeculae, and higher trabecular number, bone connectivity and SMI values, indicating a shift towards a rod-like structure. The trabeculae’s higher slenderness ratio and lower number and connectivity will affect a reduction in the trabeculae’s buckling strength ^42,43^, resulting in the vertebral body becoming more compliant. Combined with the projected shift in loading from the cancellous bone to the vertebral cortex ^29^, which has a more central role in rodent vertebral strength than in human vertebrae ^29^, both mechanisms will likely cause the observed decrement and higher variability of vertebral stiffness, resulting in the degradation of stiffness having a more dominant effect on the rodent vertebrae’s strength.

Regression analysis found that higher vertebral stiffness and BMD explained 60% of the irradiated vertebrae’s strength. Employing FE simulation, Emerzian et al. ^21^ suggested the observed degradation of the irradiated vertebral mechanical response was dominated by alteration of the bone elastic (i.e., mineral) properties. In contrast to previous longitudinal irradiation studies in mice ^27^ and rats ^21^ reporting that low levels of irradiation had little effect on TMD values, application of high dose irradiation in our study was observed to cause a marked increase in TMD at the early stages of the study, which decreased sharply at the later stages of the study, a pattern matched by the serum NTX values. Furthermore, correlation analysis at the terminal assessment revealed that, in contrast to the control animals exhibiting a strong positive correlation between TMD and vertebral strength and stiffness, the irradiated vertebrae’s strength and stiffness showed a markedly weaker correlation with TMD. In agreement with previous studies ^12,21,27,43^, we found that higher AGEs values in the irradiated bones were not associated with the reduction in measured vertebral strength. However, we observed a positive correlation between AGEs and stiffness, which also correlated with vertebral strength. This suggests that irradiation damages the bone tissue’s elastic property (stiffness), via the direct accumulation of AGEs inducing a stiffer but more brittle matrix, which resulted in increased fragility, impacting the bone’s inelastic property (strength) ^27^.

The regression model showed that lower collagen content significantly contributed to the irradiated vertebral strength and stiffness. In agreement with previous studies ^21,27,44^, our findings suggest that, beyond bone architecture, mass and volume fraction, additional factors not captured by micro-CT measures may significantly contribute to the temporal degradation in bone modulus and thus strength. Mouse hindlimb irradiation models reported that irradiated bone, three months after treatment, exhibited higher collagen crosslinks and matrix/mineral alignment, ^43^, a decreased mineral to matrix ratio ^12,18^ and reduced collagen energy-to-failure ^15^ at 3 months post-irradiation. Further studies are needed to confirm whether high radiation doses alter cellular activity and bone tissue quality, leading to changes in elastic properties and thereby weakening the structural response of irradiated vertebrae.

This study has several limitations. First, our rat model was of healthy vertebral bone. Palliative RT is clinically administered to metastatic spine patients. Thus, our study cannot realistically capture the combined effects of tumor burden and IR-induced bone damage, nor can it capture the likely effect of the irradiation on bone homeostasis due to lesion integration within the irradiated region. However, choosing a non-cancer model allows us to minimize the variance in vertebral bone properties and strength, likely to be introduced by the bone metastasis ^7,9^. Second, quadrupeds and humans differ substantially in locomotion patterns, spinal curvature, and local inter-vertebral kinematics, which produce different spinal loading. They also differ in vertebral geometry, bone microstructure, and potentially bone tissue material properties. Taken together, these differences may limit the translatability of our rat model findings to the effects of irradiation on metastatic human vertebral bone. Third, our mechanical testing was limited to the L4 vertebral level. Therefore, the observed effects of irradiation on vertebral bone properties may be site-specific, differing across spinal regions and between vertebral cancellous bone and the cancellous and cortical bone of the appendicular skeleton. Our mechanical assessment was also performed as an evaluation of strength, defined as failure under a monotonic single event. However, VF likely occurs within days to a year post-irradiation ^5^ suggesting that both time-dependent (visco-elastic) and time-varying (fatigue) based loading may influence the initiation and progression of VF. Fourth, our study employed only male rats. Emerzian et al ^21^ showed that sex influences the impact of irradiation on trabecular thickness, separation, and connectivity. However, these sex-specific microstructural differences (8% in males vs 24% in females) did not significantly affect the treatment’s overall impact on the vertebral biomechanical strength, concluding that changes in bone mass and tissue material properties were the dominant factors. Finally, our study only evaluated outcomes up to 28 days post-irradiation. Finally, our study evaluation period culminated in 28 days post-IR. With rodents having a metabolic rate four to six times higher than that of humans ^45^, this period corresponds to a follow-up period of up to six months in humans. We designed the study to capture this period due to clinical findings reporting 13.9% - 39% of cancer patients treated with SBRT suffering VF within six months post-SBRT ^5^.

In summary, our preclinical study showed that high-dose radiotherapy disrupts bone remodeling and compromises vertebral mechanical properties in a time-dependent manner. The marked divergence in the early days post-irradiation, corresponding to 2-3 months in human patients, was dominated by trabecular perforation and connectivity loss, while tissue mineralization alterations appeared later and were less pronounced. By contrast, at the later endpoint, corresponding to 4-6 months post radiotherapy in human patients, there was marked degradation in architectural indices and bone tissue composition, superseding the degree of loss in mechanical vertebral strength. Specifically, our findings regarding the contribution of bone stiffness, driven by loss of bone trabeculae and tissue density and the transition to a strut-dominated spatial architecture, suggest irradiation affects pre-yield bone properties, causing loss of vertebral strength beyond the contribution of bone mineral density. These findings may help to explain why the increase in fracture risk in patients undergoing RT shows a weak association with the increase in bone mineral density alone ^7^. These results emphasize the need for a comprehensive assessment of countermeasures to restore bone cellular homeostasis and for improved clinical measures of bone quality degradation beyond bone mass to better manage the risk of vertebral fractures in patients receiving palliative radiation.

## Acknowledgments

This research was supported by the National Institute of Arthritis and Musculoskeletal and Skin Diseases, which supported the work of R. Alkalay under its Research Project Grants (AR075964)

**Figure A.1:** Heat map highlighting significant correlations between the bone architectural indices, composition and mechanical properties in the control vertebrae at terminal assessment. Colors closer to brown indicate higher positive correlation, and colors closer to blue represent negative correlation.

## References

1 van der Velden, J. M. et al. Prospective evaluation of the relationship between mechanical stability and response to palliative radiotherapy for symptomatic spinal metastases. Oncologist 22, 972–978 (2017).

2 Huo, M. et al. Stereotactic spine radiosurgery: Review of safety and efficacy with respect to dose and fractionation. Surg Neurol Int 8, 30 (2017). 10.4103/2152-7806.200581

3 Guckenberger, M., Dahele, M., Ong, W. L. & Sahgal, A. Stereotactic Body Radiation Therapy for Spinal Metastases: Benefits and Limitations. Semin Radiat Oncol 33, 159–171 (2023). 10.1016/j.semradonc.2022.11.006

4 Pierce, S. M. et al. Long-term radiation complications following conservative surgery (CS) and radiation therapy (RT) in patients with early stage breast cancer. . Int J Radiat Oncol Biol Phys 23, 915–923 (1992).

5 Sahgal, A., Whyne, C. M., Ma, L., Larson, D. A. & Fehlings, M. G. Vertebral compression fracture after stereotactic body radiotherapy for spinal metastases. Lancet Oncol. 14, e310–320 (2013).

6 Van den Brande, R., et al. Epidemiology of spinal metastases, metastatic epidural spinal cord compression and pathologic vertebral compression fractures in patients with solid tumors: A systematic review. J Bone Oncol 35, 100446 (2022). 10.1016/j.jbo.2022.100446

7 Confavreux, C. B., Follet, H., Mitton, D., Pialat, J. B. & Clezardin, P. Fracture Risk Evaluation of Bone Metastases: A Burning Issue. Cancers (Basel) 13 (2021). 10.3390/cancers13225711

8 Gaddipati, R., Jensen, G. L., Swanson, G., Hammonds, K. & Morrow, A. The Effect of High-Dose Radiation Therapy on Healthy Vertebral Bone Density. Cureus 14, e22565 (2022). 10.7759/cureus.22565

9 Wei, R. L. et al. Bone mineral density loss in thoracic and lumbar vertebrae following radiation for abdominal cancers. Radiother Oncol 118, 430–436 (2016). 10.1016/j.radonc.2016.03.002

10 Costa, S. & Reagan, M. R. Therapeutic Irradiation: Consequences for Bone and Bone Marrow Adipose Tissue. Front Endocrinol (Lausanne) 10, 587 (2019). 10.3389/fendo.2019.00587

11 Okunieff, P. et al. Radiation-induced changes in bone perfusion and angiogenesis. Int J Radiat Oncol Biol Phys 42, 885–889 (1998). 10.1016/s0360-3016(98)00339-3

12 Bartlow, C. M., Mann, K. A., Damron, T. A. & Oest, M. E. Limited field radiation therapy results in decreased bone fracture toughness in a murine model. PLoS One 13, e0204928 (2018). 10.1371/journal.pone.0204928

13 Donaubauer, A. J. et al. The Influence of Radiation on Bone and Bone Cells-Differential Effects on Osteoclasts and Osteoblasts. Int J Mol Sci 21 (2020). 10.3390/ijms21176377

14 Wright, L. E. et al. Single-Limb Irradiation Induces Local and Systemic Bone Loss in a Murine Model. J Bone Miner Res 30, 1268–1279 (2015). 10.1002/jbmr.2458

15 Bartlow, C. M., Mann, K. A., Damron, T. A. & Oest, M. E. Altered mechanical behavior of demineralized bone following therapeutic radiation. J Orthop Res 39, 750–760 (2021). 10.1002/jor.24868

16 Oest, M. E., Policastro, C. G., Mann, K. A., Zimmerman, N. D. & Damron, T. A. Longitudinal Effects of Single Hindlimb Radiation Therapy on Bone Strength and Morphology at Local and Contralateral Sites. J Bone Miner Res 33, 99–112 (2018). 10.1002/jbmr.3289

17 Wernle, J. D., Damron, T. A., Allen, M. J. & Mann, K. A. Local irradiation alters bone morphology and increases bone fragility in a mouse model. J Biomech 43, 2738–2746 (2010). 10.1016/j.jbiomech.2010.06.017

18 Mandair, G. S. et al. Radiation-induced changes to bone composition extend beyond periosteal bone. Bone Reports 12, 100262 (2020).

19 Barth, H. D., Launey, M. E., Macdowell, A. A., Ager, J. W., 3rd & Ritchie, R. O. On the effect of X-ray irradiation on the deformation and fracture behavior of human cortical bone. Bone 46, 1475–1485 (2010). 10.1016/j.bone.2010.02.025

20 Akkus, O. & Rimnac, C. M. Fracture resistance of gamma radiation sterilized cortical bone allografts. J Orthop Res 19, 927–934 (2001). 10.1016/S0736-0266(01)00004-3

21 Emerzian, S. R. et al. Relative Effects of Radiation-Induced Changes in Bone Mass, Structure, and Tissue Material on Vertebral Strength in a Rat Model. Journal of Bone and Mineral Research 38, 1032–1042 (2023).

22 Igarashi, T. et al. Effects of radiation on the bone strength of spinal vertebrae in rats. Spine 47, E514–E520 (2022).

23 Dabbagh Ohadi, M. A., et al. Single-fraction versus multifraction stereotactic radiosurgery for spinal metastases: systematic review and meta-analysis. Neurosurg Focus 58, E16 (2025). 10.3171/2025.2.FOCUS24985

24 Karim, L. & Vashishth, D. Heterogeneous glycation of cancellous bone and its association with bone quality and fragility. PLoS One 7, e35047 (2012). 10.1371/journal.pone.0035047

25 Benjamini, Y. & Hochberg, Y. Controlling the False Discovery Rate: A Practical and Powerful Approach to Multiple Testing. Journal of the Royal Statistical Society: Series B (Methodological) 57, 289–300 (2018). 10.1111/j.2517-6161.1995.tb02031.x

26 Bandstra, E. R. et al. Long-term dose response of trabecular bone in mice to proton radiation. Radiat Res. 169, 607–614 (2008).

27 Pendleton, M. M. et al. Relations Between Bone Quantity, Microarchitecture, and Collagen Cross-links on Mechanics Following In Vivo Irradiation in Mice. JBMR Plus 5, e10545 (2021). 10.1002/jbm4.10545

28 Wu, T., Bonnheim, N. B., Pendleton, M. M., Emerzian, S. R. & Keaveny, T. M. Radiation-induced changes in load-sharing and structure-function behavior in murine lumbar vertebrae. Comput Methods Biomech Biomed Engin 27, 1278–1286 (2024). 10.1080/10255842.2023.2239415

29 Alwood, J. S. et al. Heavy ion irradiation and unloading effects on mouse lumbar vertebral microarchitecture, mechanical properties and tissue stresses. Bone 47, 248–255 (2010). 10.1016/j.bone.2010.05.004

30 Zhai, J. et al. Influence of radiation exposure pattern on the bone injury and osteoclastogenesis in a rat model. Int J Mol Med 44, 2265–2275 (2019). 10.3892/ijmm.2019.4369

31 Alam, I. et al. Identification of genes influencing skeletal phenotypes in congenic P/NP rats. Journal of Bone and Mineral Research 25, 1314–1325 (2010).

32 Jee, W. & Yao, W. Overview: animal models of osteopenia and osteoporosis. Journal of musculoskeletal & neuronal interactions 1, 193–207 (2001).

33 Banu, J., Wang, L. & Kalu, D. N. Age-related changes in bone mineral content and density in intact male F344 rats. Bone 30, 125–130 (2002). 10.1016/s8756-3282(01)00636-6

34 Govey, P. M., Zhang, Y. & Donahue, H. J. Mechanical Loading Attenuates Radiation-Induced Bone Loss in Bone Marrow Transplanted Mice. PLoS One 11, e0167673 (2016). 10.1371/journal.pone.0167673

35 Alwood, J. S. et al. Low-dose, ionizing radiation and age-related changes in skeletal microarchitecture. J Aging Res 2012, 481983 (2012). 10.1155/2012/481983

36 Zhang, J. et al. Therapeutic ionizing radiation induced bone loss: a review of in vivo and in vitro findings. Connect Tissue Res 59, 509–522 (2018). 10.1080/03008207.2018.1439482

37 Dekker, H. et al. Osteocyte Apoptosis, Bone Marrow Adiposity, and Fibrosis in the Irradiated Human Mandible. Adv Radiat Oncol 7, 100951 (2022). 10.1016/j.adro.2022.100951

38 Wang, Y. et al. Radiation induces primary osteocyte senescence phenotype and affects osteoclastogenesis in vitro. Int J Mol Med 47 (2021). 10.3892/ijmm.2021.4909

39 Alwood, J. S. et al. Ionizing Radiation Stimulates Expression of Pro-Osteoclastogenic Genes in Marrow and Skeletal Tissue. J Interferon Cytokine Res 35, 480–487 (2015). 10.1089/jir.2014.0152

40 Willey, J. S. et al. Early increase in osteoclast number in mice after whole-body irradiation with 2 Gy X rays. Radiat Res. 170, 388–392 (2008).

41 Bakar, A. A. A. et al. Systematic Review on Multilevel Analysis of Radiation Effects on Bone Microarchitecture. Biomed Res Int 2022, 9890633 (2022). 10.1155/2022/9890633

42 Keaveny, T. M. & Hayes, W. C. A 20-year perspective on the mechanical properties of trabecular bone. J Biomech Eng 115, 534–542 (1993). 10.1115/1.2895536

43 Oest, M. E. & Damron, T. A. Focal therapeutic irradiation induces an early transient increase in bone glycation. Radiat Res 181, 439–443 (2014). 10.1667/RR13451.1

44 Gong, B., Oest, M. E., Mann, K. A., Damron, T. A. & Morris, M. D. Raman spectroscopy demonstrates prolonged alteration of bone chemical composition following extremity localized irradiation. Bone 57, 252–258 (2013).

45 Agoston, D. V. How to Translate Time? The Temporal Aspect of Human and Rodent Biology. Front Neurol 8, 92 (2017). 10.3389/fneur.2017.00092

