## Supplementary figures and images for "Palliative Irradiation Affects The Degradation Of Vertebral Bone Mechanics, Architecture, and Composition: A Longitudinal In Vivo Study In A Rat Model"

### Supplemental Figure A.1

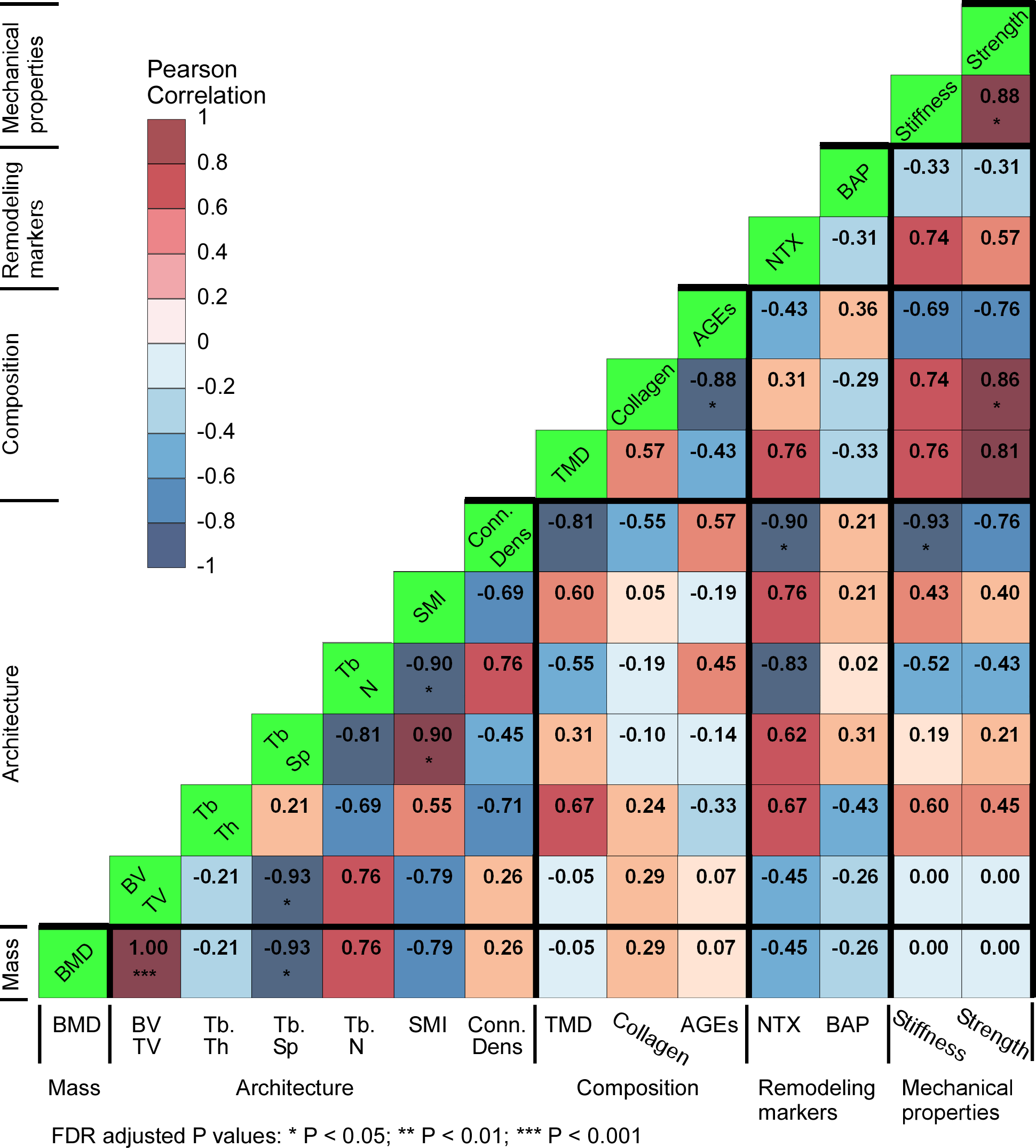
